# The effect of substrate stiffness on tensile force transduction in the epithelial monolayers

**DOI:** 10.1101/2021.09.06.459078

**Authors:** Aapo Tervonen, Sanna Korpela, Soile Nymark, Jari Hyttinen, Teemu O Ihalainen

**Affiliations:** BioMediTech, Faculty of Medicine and Health Technology, Tampere University, Tampere, Finland

## Abstract

In recent years, the importance of mechanical signaling and the cellular mechanical microenvironment in affecting cellular behavior has been widely accepted. Cells in epithelial monolayers are mechanically connected to each other and the underlying extracellular matrix (ECM), forming a highly connected mechanical system subjected to various mechanical cues from their environment, such as the ECM stiffness. Changes in the ECM stiffness have been linked to many pathologies, including tumor formation. However, our understanding of how ECM stiffness and its heterogeneities affect the transduction of mechanical forces in epithelial monolayers is lacking. To investigate this, we used a combination of experimental and computational methods. The experiments were conducted using epithelial cells cultured on an elastic substrate and applying a mechanical stimulus by moving a single cell by micromanipulation. To replicate our experiments computationally and quantify the forces transduced in the epithelium, we developed a new model that described the mechanics of both the cells and the substrate. Our model further enabled the simulations with local stiffness heterogeneities. We found the substrate stiffness to distinctly affect the force transduction as well as the cellular movement and deformation following an external force. Also, we found that local changes in the stiffness can alter the cells’ response to external forces over long distances. Our results suggest that this long-range signaling of the substrate stiffness depends on the cells’ ability to resist deformation. Furthermore, we found that the cell’s elasticity in the apico-basal direction provides a level of detachment between the apical cell-cell junctions and the basal focal adhesions. Our simulation results show potential for increased ECM stiffness, e.g. due to a tumor, to modulate mechanical signaling between cells also outside the stiff region. Furthermore, the developed model provides a good platform for future studies on the interactions between epithelial monolayers and elastic substrates.

**Author summary:** Cells can communicate using mechanical forces, which is especially important in epithelial tissues where the cells are highly connected. Also, the stiffness of the material under the cells, called the extracellular matrix, is known to affect cell behavior, and an increase in this stiffness is related to many diseases, including cancers. However, it remains unclear how the stiffness affects intercellular mechanical signaling. We studied this effect using epithelial cells cultured on synthetic deformable substrates and developed a computational model to quantify the results better. In our experiments and simulations, we moved one cell to observe how the substrate stiffness impacts the deformation of the neighboring cells and thus the force transduction between the cells. Our model also enabled us to study the effect of local stiffness changes on the force transduction. Our results showed that substrate stiffness has an apparent impact on the force transduction within the epithelial tissues. Furthermore, we found that the cells can communicate information on the local stiffness changes over long distances. Therefore, our results indicate that the cellular mechanical signaling could be affected by changes in the substrate stiffness which may have a role in the progression of diseases such as cancer.

## 1 Introduction

Our understanding of the importance of mechanical forces and microenvironment in cellular processes and signaling alongside biochemistry has drastically improved during the last decades [1–3]. Biomechanics has a vital role in embryogenesis, stem cell differentiation, tissue homeostasis, and migration [4–10]. In addition, abnormal changes in the biomechanics of the cells and the extracellular matrix (ECM) are linked to many pathological processes, including tumor- and carcinogenesis [11–15]. While the research on the effect of physical cues and the role of the mechanical environment on cell functions and signaling is active, the understanding of this mechanical system is not complete. In epithelial tissues, the cellular mechanical microenvironment is formed by the neighboring cells and the ECM on the basal side of the cells. In some instances, the apical side of the cells is subjected to shearing forces from fluid flow [16]. Thus, epithelial cells are exposed to various physical cues rising from their environment. The high interconnectivity of the epithelial tissues enables the cells to distribute exogeneous mechanical energy and transmit endogenous forces to their environment [17, 18]. Here, we investigated how the propagation of tensile forces in the epithelial monolayer is affected by the stiffness of the substrate under the cells.

Cells can use their actomyosin cytoskeleton to generate contractile forces that enable cells to change their shape and move [18, 19]. Since the actomyosin cytoskeletons of neighboring epithelial cells are connected via adherens junctions, these contractile forces can be transmitted between cells over distances [20]. Cells have various responses to these exogenous forces, behaving elastically over a short time scale by deforming and viscously over sustained stress by dissipating the stress via various processes [21–24]. Endo- and exogenous forces can also alter the structural state of different proteins, which may lead to the conversion of forces to biochemical signals via a process known as mechanotransduction [3, 25]. Furthermore, mechanical forces have been indicated to have an even more direct effect on the cell behavior since they are transmitted directly to the cell nucleus along the actin stress fibers, where they have been shown to be able to modulate gene expression [26–29]. Therefore, understanding how forces are transmitted between cells is essential to understand the mechanical system formed by the epithelial tissues.

Epithelial cells are connected to the ECM on their basal side via focal adhesions, which connect the actomyosin cytoskeleton to the basal lamina. The focal adhesions contain mechanosensitive proteins that enable the cells to sense the external forces and ECM stiffness [30–32], which has been shown to affect many cell functions during development, homeostasis, and diseases. For example, ECM stiffness has been heavily linked to cell differentiation [6, 33] and the metastatic potential of tumors [34–36]. It is well established that tumor stroma, the adjacent tissue surrounding the tumor, is often considerably stiffer than native tissue [37, 38]. In tissues, this leads to stiffness gradients and interfaces between the stroma and the surrounding healthy tissue, which can influence cellular mechanosignaling, especially during cancer invasion [14, 39]. However, we do not fully understand how the ECM stiffness or stiffness gradients affect the mechanical signaling between cells in confluent epithelial monolayers.

There is a plethora of computational methods that can describe the mechanical system formed by the epithelial monolayer. Vertex models are a relatively simplistic approach that reduces the mechanical properties of the cells to only a few parameters [40–42]. Methods such as the subcellular element method [43, 44] and immersed-boundary method [45–47] provide a more nuanced description of the cells and their mechanical properties but differ on how they describe the cells and solve the cellular movement. Only a few cell-based models have included a description of a deformable substrate under the cells [42, 43].

This study aimed to describe how strain and forces propagate in an elastic mechanical system formed by the epithelial monolayer and a deformable substrate with different stiffnesses. The work was conducted experimentally using Madin-Darby canine kidney (MDCK) II cell model on polyacrylamide (PAA) hydrogel substrates and computationally using a cell-based model we developed. To investigate the propagation of tensile forces between the neighboring cells and the cells and the ECM, we, experimentally and in the computational model, moved a single cell, causing a local stretching in the epithelium. Furthermore, we used our computational model in combination with data from the literature to study the mechanics related to subcellular changes in cell shapes.

## Results

### Micromanipulation of epithelial monolayers on substrates with varying stiffness

To experimentally study the effect of environmental stiffness on the propagation of forces and deformation in epithelial tissue, we used an *in vitro* model of MDCK II cells expressing tight junction marker mEmerald-Occludin cultured on PAA hydrogel substrates with embedded fluorescent beads. We used collagen-I-coated PAA substrates with four stiffnesses (Young’s moduli): 1.1, 4.5, 11, and 35 kPa. We manipulated a single cell with a sharp pipette attached to a piezo-driven micromanipulator as a mechanical stimulus. The pipette was brought into contact with the cell, and the micromanipulator was used to move the pipette 30 µm parallel to the surface in 1 second (speed 30 µm/s) while simultaneously imaging both the mEmerald-Occludin and the fluorescent beads.

The micromanipulation led to large deformation of the epithelium and displacement of the cells and the PAA substrate. We visualized the movement by comparing the images of the epithelium and the substrate before and following the micromanipulation (Fig. 1). It is clear from the movement of the cell boundaries (Fig. 1A) that the PAA stiffness profoundly affects the distance that the mechanical strain spreads around the manipulated cell. For example, the movement of the cell boundaries along the axis of the pipette movement (Fig. 1B, dashed lines in Fig. 1A) showed that for the 1.1-kPa substrate, the cell boundary at the edge of the imaged field (approximately 80 µm from the initial pipette position) moves 3.3 µm. This was in stark contrast with the 11-kPa and the 35-kPa substrates, for which the discernible cell boundary movement happened only at the distance of approximately 50 µm along the axis of its movement (Fig. 1B).

**Fig 1.**
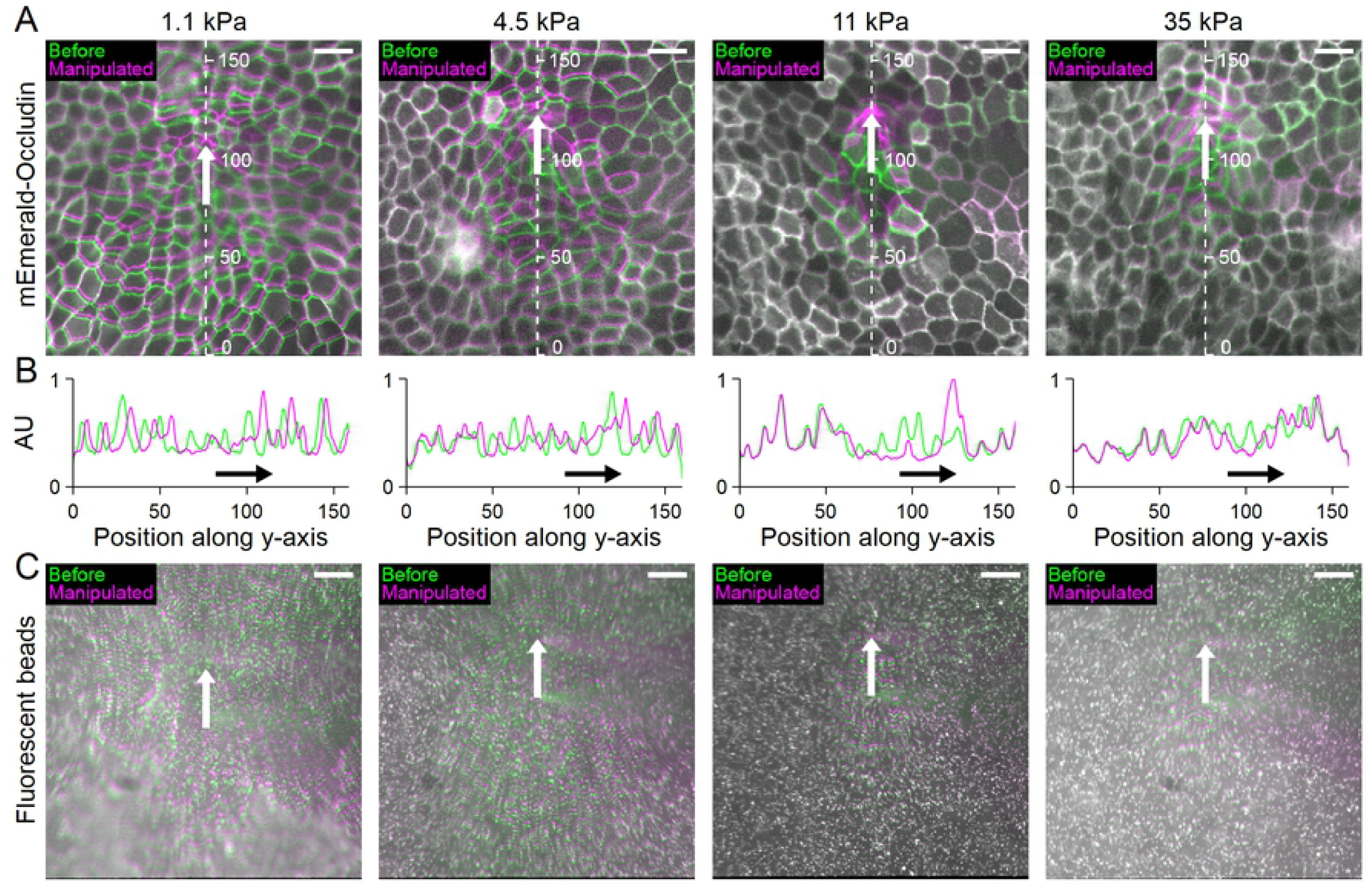
Displacements of MDCK II cells and substrate with different stiffnesses. (A) Representative examples of mEmerald-Occludin-expressing MDCK II cell movement on polyacrylamide (PAA) hydrogel substrates with stiffnesses of 1.1, 4.5, 11, and 35 kPa following the movement of the micromanipulated pipette for 30 µm in 1 s (pipette movement shown by the white arrow). The boundaries of cells were indicated using mEmerald-Occludin and shown in green before the micromanipulation and in magenta following the 30 µm pipette movement. The cell displacement on the right side of the pipette (white arrow) is partly due to the pipette affecting part of the image. (B) Line plot along the dashed lines for cell boundaries in A in arbitrary units (AU) before (green) and following the micromanipulation (magenta) for each gel stiffness to better show the magnitude of the cell movement along the pipette movement axis. The data was smoothed using 10 pixel moving average. The pipette movement is indicated by the black arrows. (C) The movement of the fluorescent beads embedded in the PAA hydrogel substrates with the corresponding stiffness underlying the epithelia shown in A. The pipette movement of 30 µm is shown by the white arrow and the bead locations before and following the micromanipulation in green and magenta, respectively. The pipette shadow was affecting the results on the right side of the white arrow. Scale bar, 20 *µ*m.

Similar to the cells, the displacement of the substrate was naturally affected by their stiffness. There was deformation in the whole imaged field for both the 1.1- and 4.5-kPa substrates (Fig. 1C) in the direction parallel to the pipette movement. However, the displacement in the perpendicular direction was more limited with the 4.5-kPa substrate than with 1.1 kPa. The deformation was even smaller with the stiffer (11 and 35 kPa) substrates (Fig. 1C).

To quantify the cell displacements, we segmented the cell imaging data before and following the micromanipulation to obtain the cell outlines. Using the outlines, we defined a geometrical cell center, which we then used to measure the displacement of the individual cell during the micromanipulation. This provided us with a spatial map of the cell center movements in relation to their distance and direction from the initial position of the pipette (*p*_*0*_). We then interpolated the cell center movement data over the whole imaging area to obtain a continuous distribution for each measurement and calculated the average distribution for each substrate stiffness (Fig. 2A and B). In order to do the same for the substrate data, we used particle image velocimetry (PIV) analysis to find the displacement of the substrate beads between images taken before and following the micromanipulation. Similar to the cell data, this was averaged and plotted in relation to *p*_*0*_ (Fig. 2C and D).

**Fig 2.**
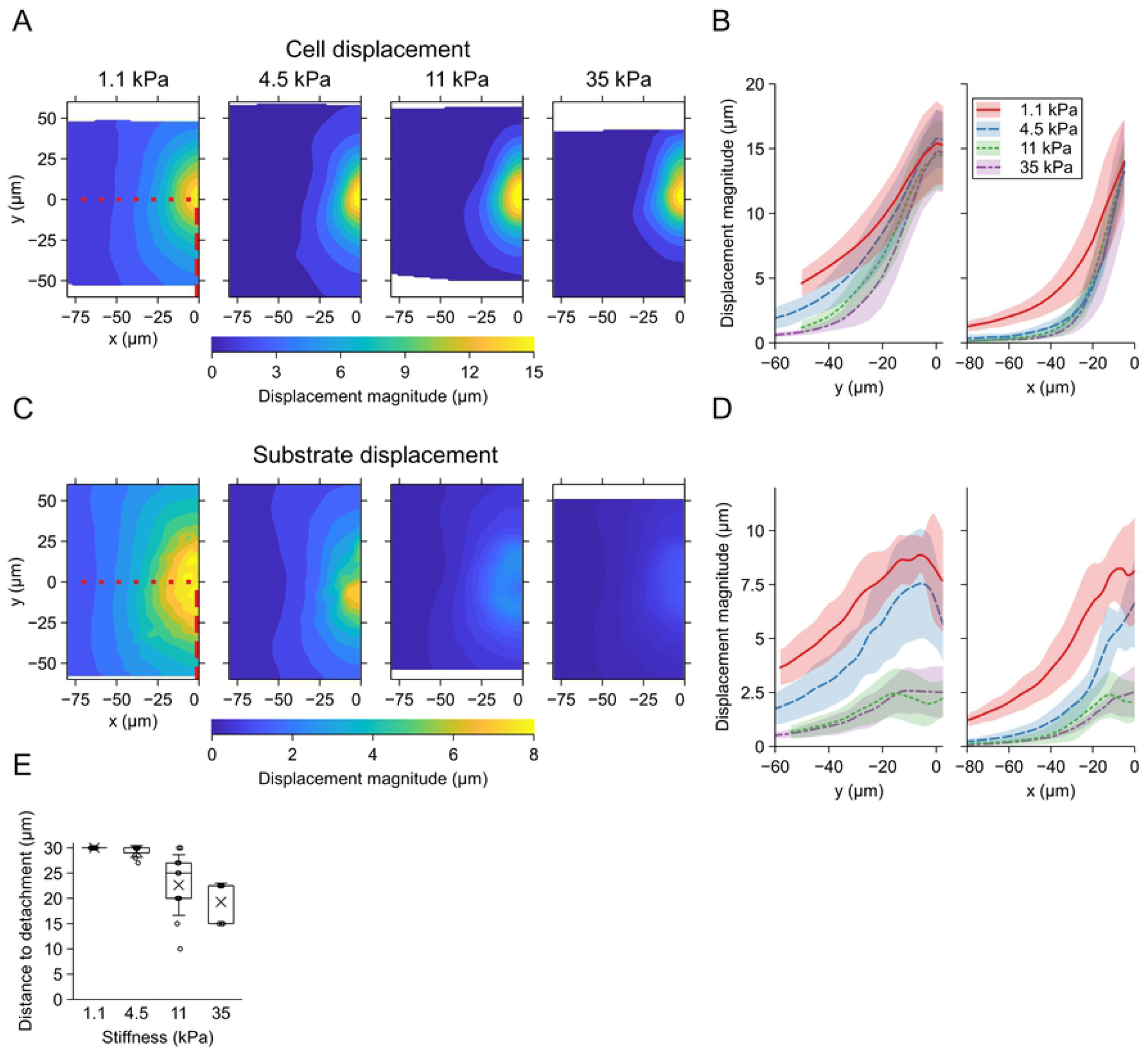
Displacement fields of the MDCK II cells and polyacrylamide (PAA) hydrogel substrates. (A) Average displacement of the segmented MDCK II cell centers as function of location of the original cell positions in relation to the initial pipette position (p_0_) for the stiffnesses 1.1 (n = 11), 4.5 (n = 7), 11 (n = 11), and 35 kPa (n = 7). The field is limited to the left of micromanipulation axis since the movement was symmetric on either side of the axis. The area of the shown displacement field varies between the stiffnesses since *p*_0_ in relation to the imaging area varied between measurements. (B) The cell center displacement along the y-axis (red dashed line in A) away from the direction of the pipette movement (left) and along x-axis (red dotted line in A) perpendicular to the pipette movement direction (right) for each stiffness. (C) Average displacement of PAA hydrogel substrates based on the particle image velocimetry (PIV) analysis as function of location in relation to *p*_0_ for the stiffnesses 1.1 (n = 11), 4.5 (n = 7), 11 (n = 11), and 35 kPa (n = 7). The field is limited to the left of micromanipulation axis since the movement was symmetric on either side of the axis and the pipette causes artefacts in the PIV data on the right side of the pipette. The area of the shown displacement field varies between the stiffnesses since the pipette position in relation to the imaging area varied between measurements. Note that maximum displacement is different than with the cells. (D) The PAA substrate displacement along the red dashed line in C away from the direction of the pipette movement (left) and along the red dotted line in C perpendicular to the pipette movement direction (right) for each stiffness. The shaded region represents the SD for each stiffness. (E) Distance of pipette movement before cells detached from the substrate estimated from the live imaging data for the different stiffnesses 1.1 (n = 11), 4.5 (n = 7), 11 (n = 11), and 35 kPa (n = 7). The indicated cases with the distance to detachment of 30 µm did not detach from the substrate during the experiment.

Interestingly, the 30-*µ*m pipette movement translated to a cell center movement of a similar range independent of the substrate stiffness with values of 15.4*±* 3.2, *±*15.8 2.3, *±* 14.5 *±*2.6, *±* and 14.8 *±* 3.1μm (mean *±* SD), from the softest to the stiffest substrate. This difference between the pipette movement and the cell displacement can be explained by the deformation and stretching of the manipulated cell. The pipette movement caused substantial deformation to the adjacent cells in the direction of the movement, and thus the displacement of these cells was difficult to quantify. Therefore, we mainly concentrated our analysis on the area where the pipette pulled and stretched the cells and present the results mainly as a function of the location on the negative y-axis from *p*_*0*_. Parallel to the pipette movement (Fig. 2B, along the red, dashed line in 2A), the cell centers move 5 *µ*m or more within a distance of 47, 34, 26, and 20 *µ*m from *p*_*0*_ respectively for 1.1, 4.5, 11, and 35-kPa substrates. Interestingly, the three stiffest substrates have the same amount of cell center displacement perpendicular to the pipette movement. In contrast, with the softest substrate, the displacement of at least 5 *µ*m extends to approximately 1.5 times farther away than the rest (Fig. 2B).

The amount of displacement of the substrate was considerably smaller than that of the cells (Fig. 2D). The maximum displacements, located near *p*_*0*_, were, from the softest to the stiffest, 8.9*±* 0.8, 7.6*±* 2.5, 2.7*±* 1.0, and 2.6*±* 1.1 *µ*m. Therefore, the relative magnitude of the maximum substrate displacement compared to that of the cells corresponding to the stiffnesses from 1.1 to 35 kPa were 0.58, 0.48, 0.19, and 0.17. This difference in the maximum displacements partly originated from the facts that the cell-cell junctions are near the cells’ apical surface and the substrate-binding focal adhesions are on the basal side, and that the cell height provides some elasticity. The measured substrate displacements close to *p*_*0*_ were affected in the PIV analysis by the shadow of the pipette and a slight out-of-focus indentation caused by the pipette pushing the cells.

We also observed cells around the pipette detaching from the substrate in 28 % of the measurements with the 4.5-kPa substrate, in almost all the cases for the 11-kPa substrate, and in all the cases for the stiffest 35-kPa substrate (Fig. 2E and S3 Video). The detachments occurred late in the movement on the 4.5-kPa substrate, whereas those for the two stiffer substrates occurred much earlier. In addition, interestingly, there was more variance in the detachment distance for the 11-kPa substrate compared to the others. The detachment with the two stiffer substrates explained the minuscule difference between the substrate displacements in these measurements.

### Computational modeling of force propagation in the epithelium

In order to further understand the force propagation in the epithelial monolayers, we developed a cell-based computational model of the epithelial cell-cell and cell-substrate force transduction. The cells were represented by closed polygons, and the model was evolved by calculating forces between the polygon vertices similar to Tamulonis et al. [48]. We described the deformable substrate under the cells as a triangular grid of points whose movement was solved similar to that of the cell vertices. For a detailed explanation of the model, the fitting, and the simulations, see the description in S2 Text. We used the experimental results from our *in vitro* cell model and data from the literature to fit the model parameters. The computational model was first used to grow virtual epithelia, followed by the simulation of single-cell mechanical manipulation. During the simulated manipulation, we restricted the remodeling of the cell properties to describe the purely elastic properties of the experimental time scale.

We fitted the model parameters by comparing the cell center and substrate displacements between the *in vitro* experiments and the computational model. Due to the similarity of the experimental 11 and 35-kPa results in cellular displacements (Fig. 2B and D) and detachment distances (Fig. 2E), we decided to omit the 35-kPa substrate from our simulations. We assumed that the elastic properties of the epithelium are similar between the substrates, and therefore we used the same cell parameter values for each substrate stiffness in the fitting process. However, the only exception was the focal adhesion strength parameter, which we assumed to depend on the substrate stiffness and was thus altered accordingly. Similar to the experiments, in the simulation data analysis, we focused on the area of the epithelium under tension (i.e., y < 0). The most drastic movement of the cell boundaries during the micromanipulation was visible for the softest substrate (Fig. 3A). We measured the average cell and substrate displacements, which show highly similar behavior to the experiments (Fig. 3C-E).

**Fig 3.**
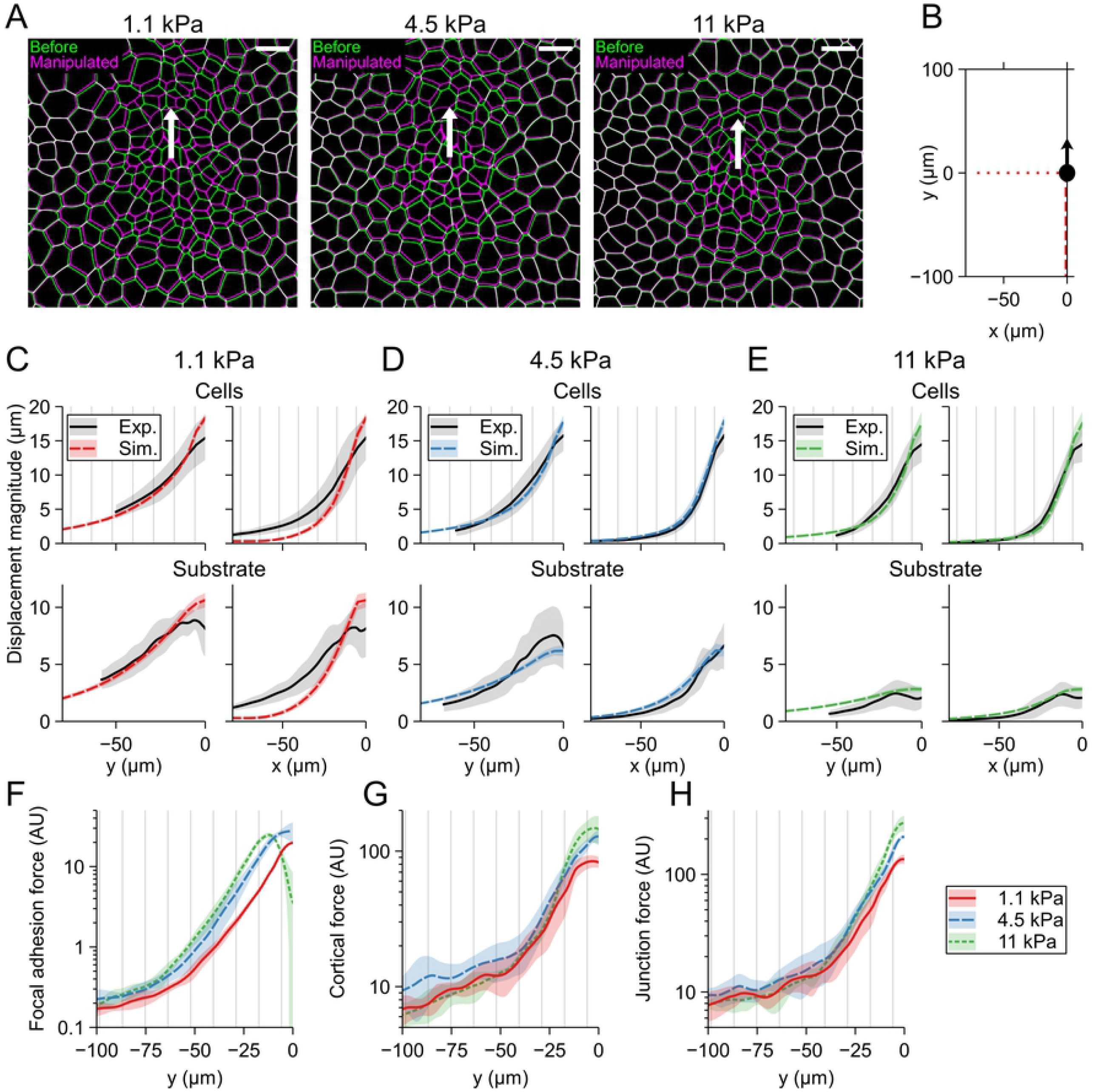
Computational model fitting results. (A) Representative figures showing the cell displacement in the simulations during the micromanipulation for the 1.1, 4.5, and 11-kPa substrates highlighting the cell shapes before (green) and following the micromanipulation (magenta). The white arrow indicates the micromanipulator movement. The scale bars are 20 *µ*m. (B) Description of the axis the results were plotted on in C-H. Comparison between the experimental cell and substrate displacement with the fitted computational model for (C) 1.1, (D) 4.5, and (E) 11-kPa substrates. The top row for each stiffness shows the fit for cell displacement in vertical (left, dashed line in B) and horizontal (right, dotted line in B) directions and the bottom row shows the same for the substrate displacements. (F) The focal adhesion, (G) the cortical, and (H) the junction forces parallel the pipette movement (dashed line in A) for the cells on 1.1, 4.5, and 11-kPa substrates in arbitrary units (AU). The force magnitudes are comparable between each other. The shaded region represents the SD. For each set of simulation parameters, n = 5.

The values obtained for the focal adhesion strengths were 0.5, 0.8, and 1.0 g s^−2^ *µ*m^−2^ for 1.1, 4.5, and 11-kPa substrates, respectively. Since we described the focal adhesions by springs, the units include the unit of the spring constant (g s^−2^). In addition, the strength depends on the length of the membrane (*µ*m) that each focal adhesion spring represents. Furthermore, it is essential to note that since the cell polygons represent the apical surface of the cells, the focal adhesion springs that connect the apical polygon vertices to the basal substrate also include the cell elasticity in the apico-basal axis. However, the increase in the focal adhesion strength as a function of substrate stiffness still reflects the stronger binding of the cell to a stiff substrate.

We analyzed the computational cell displacements similar to the experimental results by calculating the average cell center movement distributions in relation to *p*_*0*_. The substrate displacement, on the other hand, was defined directly from the substrate point movement. We transformed the results so that they were in relation to p_0_ and averaged over multiple simulations. The fitted model captured extremely well the general behavior of the *in vitro* micromanipulation (Fig. 3A), especially in the region parallel to the pipette movement (left side plots of Fig. 3C-E). However, the model produced higher cell displacement near *p*_*0*_ for all stiffnesses, most likely due to the difficulty of cell segmentation in the *in vitro* data in this area. Also, since these areas were affected in the PIV analysis, the substrate displacements here differ between the experiments and the model. The model was unable to accurately describe the displacement perpendicular to the pipette movement direction with the 1.1-kPa substrate. Furthermore, the substrate deformations remained higher over longer distances for the 4.5 and 11-kPa substrates in the model for both the cells and the substrate (left side plots in Fig. 3D and E). This may indicate that cellular structures near the pipette are damaged, therefore dampening the transmission of forces between the cells. Interestingly, the variabilities in the displacement between the computationally manipulated epithelia were small even though we ran each simulation using a different epithelial system.

The model was then used to compare the propagation of different forces depending on the substrate stiffness. To describe the forces following the micromanipulation as a function of position in relation to the *p*_*0*_, we averaged the forces over the vertices of each cell and assigned them to the original cell center positions. Next, we interpolated the averaged force magnitudes between the cell centers to obtain continuous spatial distributions and then averaged over multiple simulations.

We concentrated on the forces that arise from the interactions between the cells and the substrate (focal adhesion forces, Fig. 3F), the mechanical tension and deformation of the cell itself (cortical force, Fig. 3G), and the mechanical connection between the cells (junction forces, Fig. 3H). The substrate stiffness had an apparent effect on the maximum magnitudes of each of the three forces. The average maximum focal adhesion forces near *p*_*0*_ were 19.7, 27.6, and 24.6 AU (arbitrary units) from the softest 1.1 to the stiffest 11 kPa substrate (Fig. 3F). The maximum value for the 11-kPa substrate is lower due to the cell detachment near *p*_*0*_. The corresponding average maximum values for the cortical forces were 83.5, 127.7, and 152.1 AU (Fig. 3G) and for the junction forces 133.2, 208.2, and 272.5 AU (Fig. 3H). There were only minor differences in the junction forces between the two stiffest substrates beyond the distance of 25 *µ*m from *p*_*0*_. Interestingly, the cortical force became highest for the 4.5-kPa substrate farther than approximately 50 *µ*m away. Both the junction and cortical forces remained marginally lower for the softest 1.1-kPa substrate compared to the stiffer substrates until approximately the distance of 50 *µ*m. On the other hand, the focal adhesion forces showed an apparent effect of the stiffness over the whole range of the simulated distance. For example, the focal adhesion force level sensed by the cells on 1.1-kPa substrate at 25 *µ*m from *p*_*0*_ extended on average to 40 and 47 *µ*m for 4.5 and 11 kPa, respectively (Fig. 3F).

The results indicate that the stiffness of the substrate had an apparent effect on the propagation of cell movement following an external tensile force. However, while the forces between cells and cell deformation in the apical plane, as indicated by the increased cortical force, were higher near the manipulated cell on stiffer substrates, there were only minor differences in the forces at longer distances. On the other hand, the focal adhesion forces on the softest substrate remained lower over the simulated distance.

### Propagation of cell-cell and cell-substrate forces over substrate stiffness gradients

Next, we investigated how stiffness gradients – describing those between stiff tumorous tissues and healthy soft tissue – affect the transduction of tensile forces between the cells. Our computational model enabled the generation of epithelial monolayers attached to substrates with stiffness gradients with different slopes. We first concentrated on studying strain propagation from a soft substrate to an area of a stiff substrate. To do this, we simulated the pipette micromanipulations with substrates that included one of the following three different types of stiffness gradients between 1.1 and 11 kPa: a stiffness interface (change in stiffness in 2 *µ*m, Fig. 4H), a sharp or a shallow gradient (changes in stiffness in 10 *µ*m or 50 *µ*m, respectively, Fig. B in S1 Appendix). The pipette was moved only 20 *µ*m in these simulations to minimize the cell detachment from the substrate. We also simulated the 20-*µ*m micromanipulations with the uniform stiffnesses of 1.1 and 11 kPa for comparison (Fig. A and B in S1 Appendix). We analyzed the results by calculating how the cell and substrate displacements and the focal adhesion, cortical, and junction forces changed compared to the situation with a uniform 1.1-kPa substrate under the cells. Therefore, we calculated how much the displacements and forces changed by comparing results from simulations with a stiffness gradient against those with a uniform stiffness. This absolute difference was defined by subtracting the latter from the first at each distance from *p*_*0*_. We also calculated the relative difference between the two cases for the forces by dividing the results with the stiffness changes by those with the uniform stiffness at each distance. An example visualizes these steps in Fig. 4A. We used the relative differences for the forces to visualize better the large relative changes in the forces independent of magnitude, especially the focal adhesion forces, that are not as clearly shown with an absolute difference.

**Fig 4.**
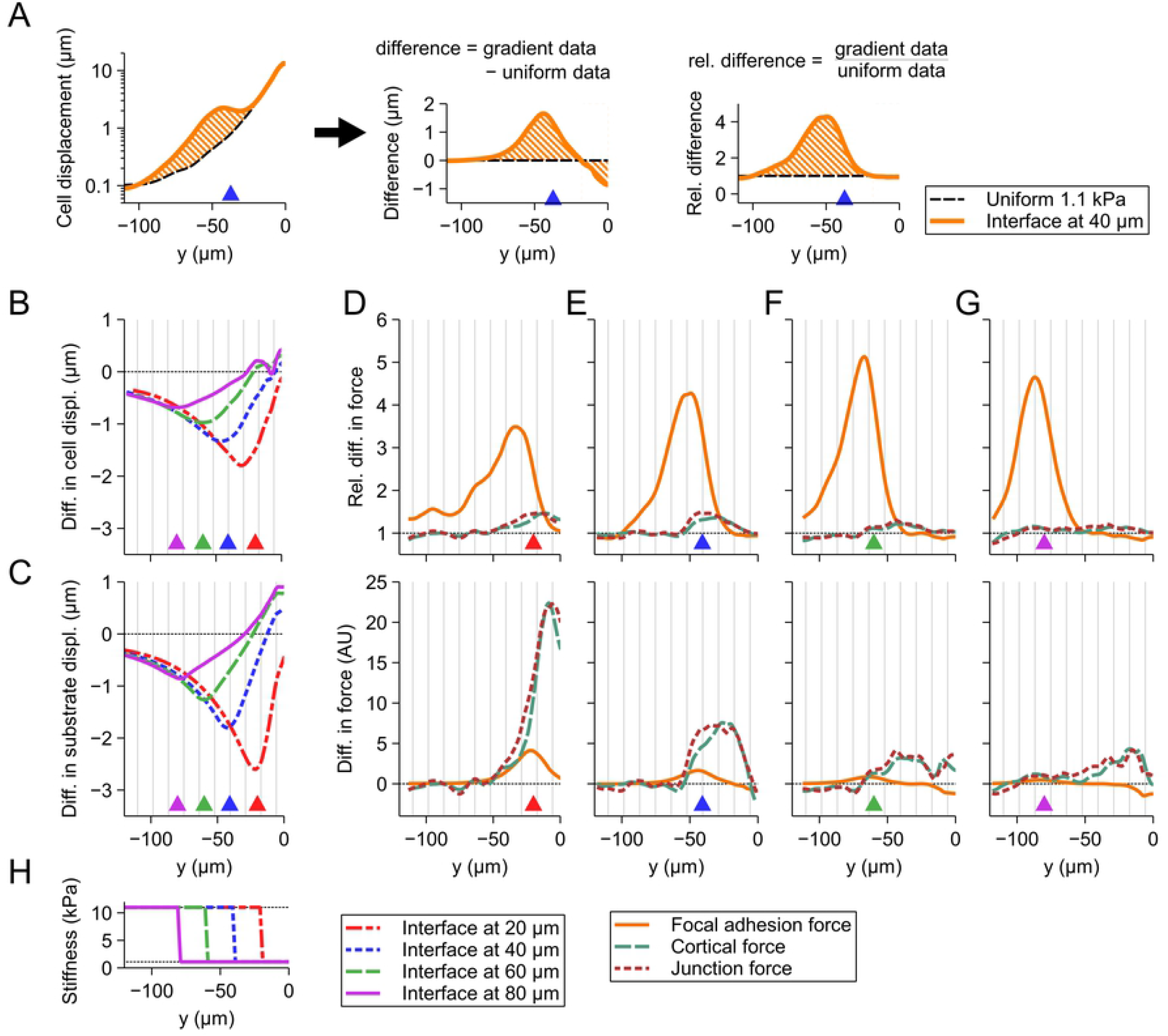
Differences in vertical cell displacement and focal adhesion, cortical, and junction force propagations caused by an interface gradient from soft to stiff substrate. The results are calculated towards the negative y-direction from the initial pipette position (*p*_0_) based on their average difference compared to the case where the substrate stiffness was the same as below the manipulated cell. (A) Example of the results calculations’ using the focal adhesion force for the stiffness interfaces at y = −40 *µ*m (blue dashed line in H). The orange striped areas correspond to each other in the figure. The difference is either calculated as absolute difference (by subtracting the results with uniform substrate from those with a gradient at each distance from *p*_0_) or as relative difference (by dividing the results with a gradient by those with the uniform stiffness at each distance from *p*_0_). The absolute difference in (B) cell and (C) substrate displacement compared to the uniform 1.1-kPa displacement for stiffness interfaces at 20, 40, 60, and 80 *µ*m. The relative (top) and absolute (bottom) differences in the average focal adhesion, cortical, and junction forces for the stiffness interfaces at (D) 20, (E) 40, (F) 60, or (G) 80 *µ*m compared to the forces in the corresponding position with 1.1-kPa substrate. (H) The stiffness interfaces for displacement and forces shown in B-G. The magnitudes of the displacement and forces are shown in Fig. A in S1 Appendix. The vertical striping shows the positions of cell boundaries for average sized cells and the positions of the interfaces are shown with the arrowheads of corresponding colors at the bottom of each figure. For each set of parameters, n = 15. AU, arbitrary unit.

Compared to the 1.1-kPa uniform substrate, the rapid increase in stiffness at the various distances from *p*_*0*_ led to a reduced cell displacement with the most prominent effect near the stiffness interface (Fig. 4B). While larger when the interface was closer to *p*_*0*_, the reduction still occurred even when the interface was up to 80 *µ*m away. However, with the interface farther away, the difference in displacement remained smaller than 1 *µ*m. The difference in the substrate displacement was slightly higher but otherwise similar to that of the cells (Fig. 4C). Interestingly, the substrate was displaced more at *p*_*0*_ when the stiffness interface was farther away.

The decreased cell and substrate displacement was accompanied by increased forces (Fig. 4D-G). The cortical and junction forces were generally increased between *p*_*0*_ and the interface and some distance beyond the interface when closer to *p*_*0*_. However, only minor changes were visible near the interface at 80 *µ*m away. On the other hand, the focal adhesion forces showed no differences between *p*_*0*_ and about one cell layer (indicated by the vertical striping) before the interface. However, these forces were greatly increased around the interface, as indicated by the peaks that continue for 3-5 cell layers on the stiff substrate. These difference peaks were most prominent in absolute terms when the interface was 20 *µ*m from *p*_*0*_ (Fig. 4D) and in relative terms when the interface at 60 *µ*m with an over 5-fold increase in the force compared to the uniform 1.1-kPa substrate (Fig. 4F). While the higher focal adhesion strengths can partly explain the increase following the interface on the stiffer substrate, the peak values were also higher than the focal adhesion forces at this distance on a uniform 11-kPa substrate (Fig. A in S1 Appendix). Our results suggest that cells situated on a soft island move less and are subjected to larger cortical and junction forces if one cell experiences a substantial deformation or movement. In addition, the cells near the stiffness interface sense high focal adhesion forces.

The observed behavior was similar with the shallow and sharp stiffness gradients compared to the interface gradients in equal distances from p_0_ (Fig. B in S1 Appendix). These gradients also produced similar relative peaks in the focal adhesion forces; however, the wider the stiffness gradient was, the more spread out and lower the peak was.

Next, we wanted to investigate how the force transduction is altered when the substrate stiffness gradient is oriented from stiff to soft. Thus, we simulated the 20-*µ*m micromanipulation of a single cell with the stiffness profiles mirroring those in the previous section. Here, the cell and substrate movements were considerably increased in comparison to uniform 11-kPa substrate (Fig. 5, Fig. D in S1 Appendix). The cell displacements were increased at the stiffness interface gradients (Fig. 5A), with the largest difference with the interfaces closer to p_0_. Interestingly, the displacement was increased well before the interface itself, also for those farther away from p_0_. The general behavior of the difference in the substrate displacement was similar to that of the cells but with slightly higher peak values (Fig. 5B). The substrate displacement was also increased at p_0_, unlike with the cells.

**Fig 5.**
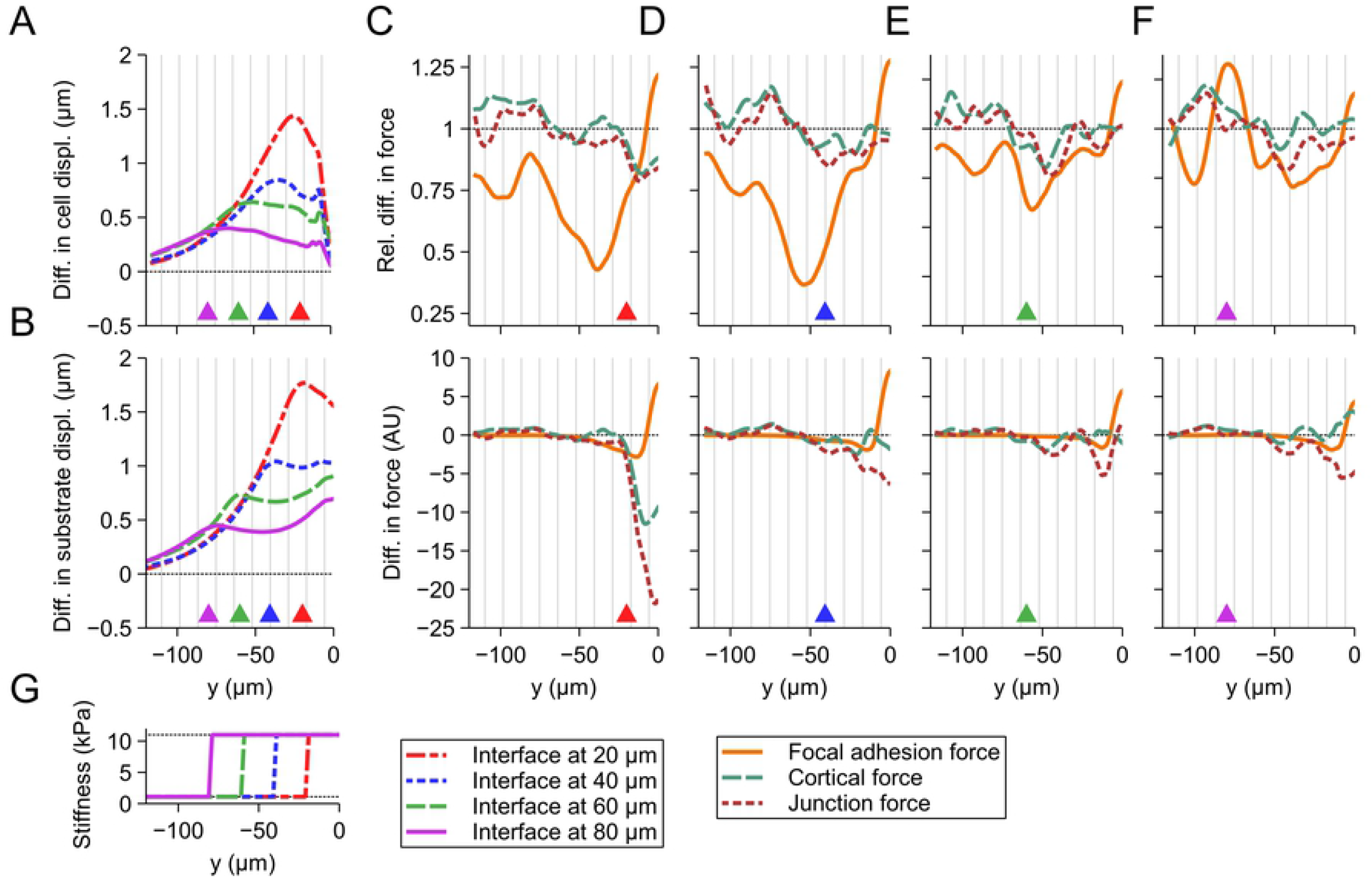
Differences in vertical cell displacement and focal adhesion, cortical, and junction force propagations caused by an interface gradient from stiff to soft substrate. The absolute difference in (A) cell and (B) substrate displacement compared to the uniform 11-kPa displacement for stiffness interface gradients at 20, 40, 60, and 80 *µ*m. The relative (top) and absolute (bottom) differences in focal adhesion, cortical, and junction forces for the stiffness interfaces at (C) 20, (D) 40, (E) 60, or (F) 80 *µ*m compared to the average cell forces in the corresponding position with 11-kPa substrate. (G) The stiffness interfaces for displacement and forces shown in A-F. The magnitudes of the displacement and forces are shown in Fig. C in S1 Appendix. The vertical striping shows the positions of cell boundaries for average sized cells and the positions of the interfaces are shown with the arrowheads of corresponding colors at the bottom of each figure. For each set of parameters, n = 15. AU, arbitrary unit. See Fig. 4A for explanation of how the absolute and relative differences were calculated.

In terms of absolute values, there are only minor differences in the forces farther than approximately 50 *µ*m from p_0_ compared to the uniform case independent of the interface gradient location (Fig. 5C-F). The cortical and junction forces were decreased close to p_0_ with the interface at 20 *µ*m (Fig. 5C). On the other hand, focal adhesion forces were increased near p_0_ but showed generally reduced values at longer distances. Furthermore, following the interfaces at 20 and 40 *µ*m, there were inverse peaks in these forces compared to the uniform substrate (Fig. 5C and D), which were not as clearly visible with the farther interfaces. The data indicates that the single-cell movement within a stiff substrate island causes larger deformations in the neighboring cells than on a uniform stiffness. Thus, the changes in the stiffness can be sensed farther away.

Again, the differences in the cell displacements and forces were similar with the shallow and sharp gradients compared to the interface gradients in similar locations (Fig. D in S1 Appendix). The reduced focal adhesion forces were also visible following the decrease in stiffness similar to those in Fig. 5C and D.

### The effect of substrate stiffness on small changes in cell shapes

Our computational model indicated that the substrate stiffness and especially stiffness gradients influence the strain distribution after a large single-cell movement. Next, we wanted to investigate how minute changes in cell-cell junctions are transmitted to the surrounding substrate. To correlate the simulations to experimental data, we simulated an optogenetic experiment, where actomyosin contractility is increased by light activation. Therefore, we implemented the optogenetic activation into our model based on the experimental and theoretical work by Staddon et al. [24]. We obtained the model parameters either directly from Staddon et al. or by fitting as described in S2 Text. We did not consider the strain-based remodeling of the cortical tension since we wanted to concentrate on the effect of the substrate stiffness on the local movement of cell boundaries, and the tension remodeling primarily affects the reduced junction length following the optogenetic activation [24]. We also allowed the remodeling of the cell structures in these simulations due to the long experiment duration compared to the micromanipulation. We increased the contractility of cell vertices forming the junctions between two cells to observe how the length of this junction reduced following the activation (Fig. 6A). We ran the simulations on substrates with uniform stiffnesses of 1.1, 4.5, or 11 kPa.

**Fig 6.**
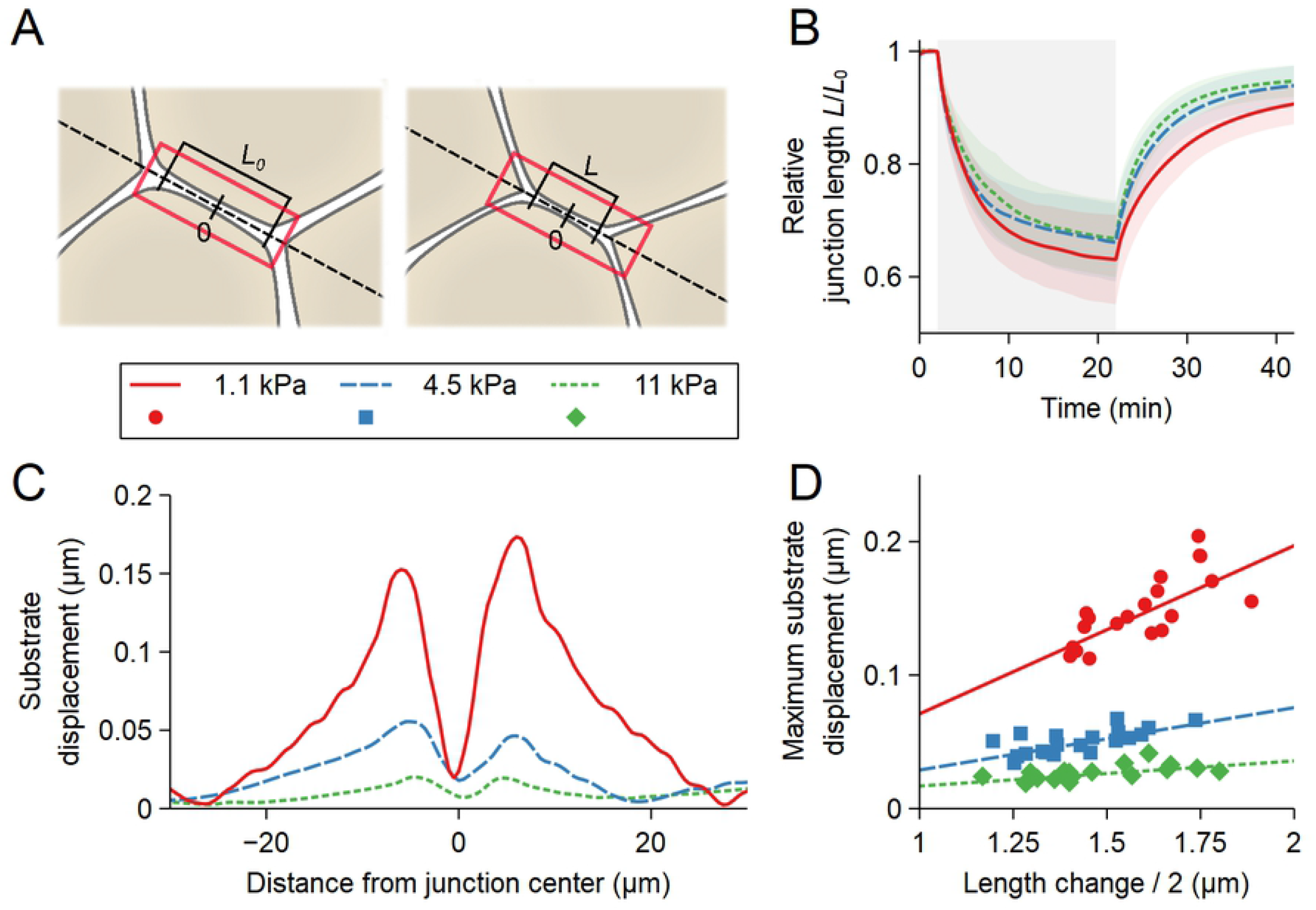
Reduction of the cell-cell junction length and substrate displacement during optogenetic activation. (A) To simulate the increased cortical contractility due to light activation, a rectangular area enclosing the cell-cell junctions between two cells (the red rectangle) was selected for activation. We then monitored the change in the junction length (L) from the initial state (L_0_) as a function of time. (B) We used a single activation of 20 minutes (grey area between 2 and 22 minutes) and calculated the relative junction length (L/L_0_) during this activation and the following relaxation. This was done for the stiffnesses 1.1, 4.5, and 11 kPa. For each stiffness, n = 20. The shaded area indicates standard deviation. (C) Representative plots of the displacement of the substrate with the three stiffnesses along the axis of the junctions (the dashed line in A) in relation to the junction center point (indicated in A). (D) Maximum substrate displacement as a function of half of the change in the junction length for each simulation for the three stiffnesses. The maximum substrate displacement was calculated as mean of the peaks on each side of the junction center point shown in C. The lines show linear fit for each set of points.

During the simulated 20-min activation, the increased cortical contractility reduced the junction length the most with the softest 1.1-kPa substrate with the final relative length of around 0.63*±* 0.08 (mean *±*SD) (Fig. 6B). The relative junction length was reduced to similar values for the two stiffer substrates with the values of 0.66 *±*0.07 and 0.67 *±*0.07 for 4.5 and 11 kPa, respectively (Fig. 6B). However, the initial length reduction was faster with the 4.5-kPa substrate than the 11-kPa, with a similar slope to the 1.1-kPa substrate.

Next, we studied how the shortening of the cell-cell junction deformed the substrate under the cells via the focal adhesions. We defined the maximum displacement of the substrate field between the moment before the activation (time = 2 min) and the end of the activation (time = 22 min) along a line defined by the activated section of junctions between two cells (the dashed line in Fig. 6A) for each simulation. Fig. 6C shows a representative displacement plot for each stiffness centered on the junction center point shown in Fig. 6A. To quantify the results, we took the average displacement peak values on each side of the junction center point. We plotted them as a function of half of the change in the junction length between the two time points for each simulation (Fig. 6D). Half of the length change was used since one of the displacement peaks in Fig. 6B resulted from half of the total junction length reduction. To compare these relative displacements with those from the micromanipulations, we calculated the mean maximum substrate displacement in relation to the corresponding half of the junction length reductions. The obtained values were 0.093, 0.035, and 0.018 respectively for 1.1, 4.5, and 11-kPa substrates, which were considerably smaller than the corresponding values in the 30-*µ*m micromanipulations, indicating a nonlinear relationship between the cell and substrate displacements.

The simulation results show that the substrate stiffness has only a minor direct effect on the small cellular morphological changes. In addition, these small changes in the cell morphology could not deform the substrates at a visible level, especially with the higher stiffnesses.

## Discussion

The role of mechanical forces in cellular communication and in the regulation of cell functions has been widely accepted [1,2, 19,29, 49]. Moreover, the stiffness of the cellular microenvironment is known to affect the cells’ mechanical properties and behavior, and an increase in the stiffness has been linked to many diseases. Most notably, in tumor formation and cancer progression, the ECM stiffness increases [11, 15, 34, 50–52], affecting how mechanical strains are transmitted between the cells. Tightly packed epithelial monolayers on deformable substrates form an excellent platform for studying how forces propagate between cells and what is the effect of the substrate stiffness in this process. To this end, we developed a computational model to describe the propagation of forces in the cell monolayer on deformable substrates based on our own experimental data and that from the literature.

We first studied how a substrate with a uniform stiffness affected the propagation of cell displacements and strain, and therefore forces, following an exogenous 30-*µ*m movement of a single cell within one second. Logically, both the cell and the substrate displacement spanned over longer distances with soft substrates. The higher substrate stiffnesses, on the other hand, greatly reduced both displacements. The high cell displacement perpendicular to the pipette movement with the soft 1.1-kPa substrate can be explained by the reported stiffnesses of the MDCK monolayers, that are between 1 and 5 kPa, when measured with atomic force microscopy [53–55]. This means that the stiffness of the epithelial monolayer in this system has a similar or slightly higher stiffness than the softest substrate and, therefore, can more readily displace the substrate than the monolayers on the stiffer substrates. We observed only minor differences between the cell displacements on the stiffer substrates, suggesting that the propagation of forces began to saturate. Therefore, having a substrate stiffer than 35 kPa would most likely not have a further effect on the cell displacements.

The difference between the maximum cell and substrate displacements can be explained by the different displacements of the apical and basal surfaces of the cell. Both our imaging of the mEmerald-Occludin-expressing MDCK cells and our computational model describe the cells by their apical surface. The displacement of the basal substrate-binding surface is more related to that of the substrate. This suggests that the cell shape in the apico-basal axis is heavily distorted, especially near the micromanipulated cell. Furthermore, the variabilities in the displacements of both the cells and the substrate predicted by our computational model were considerably smaller than those seen in the experimental results. Therefore, the variability in the epithelial morphology – i.e., the cell sizes and shapes – was not enough to explain the experimental displacement variability, and our simulation results thus only reflect an average epithelium. In reality, the mechanical properties are more varied.

We used our computational model to study the cortical, cell-cell, and cell-substrate forces during the micromanipulations. It is noteworthy to mention that the focal adhesion forces describe both the tension in the focal adhesions and the apico-basal axis of the cell. We found that all of these forces were increased by the increase in substrate stiffness. This was expected because the cells were subjected to the same external strain on the stiffer, less deformable substrates as those on the soft substrates. Therefore, the smaller substrate deformation meant that the cells were subjected to a larger portion of this strain, which led to larger cell deformation in the apical plane and higher cell strain in the apico-basal cells axis. These changes corresponded to the increases in the cortical and focal adhesion forces. Importantly, for the apical cell deformation to occur, the next cell opposite to the incoming strain must resist movement or deformation. If the next cell can be readily moved, less of the mechanical energy goes to cell deformation since it is easier to transmit onward. Therefore, the cortical and junction forces depend on each other since higher resistance against deformation leads to higher junction forces as less of the mechanical energy is absorbed by the cell cortex. However, the differences in the cortical and junction forces between the stiffnesses disappeared beyond the distance of 50 *µ*m, indicating that the bulk of the mechanical energy is absorbed closer to p_0_ on the stiff substrates.

The focal adhesion forces remained higher for the cells on stiffer substrates due to the more considerable difference between the cell and substrate displacement and the higher focal adhesion strength. Similar results were found by Goodwin et al. [56] in the developing *Drosophila* embryos, as they showed that the amount, and thus the strength, of basal cell-ECM adhesions was inversely correlated with the displacement of the apical surface. They hypothesized that the increased apical displacement was the result of more efficient apical force transmission. On the other hand, our results indicated higher forces transmitted between cells with stronger focal adhesions and smaller apical displacements in an elastic system with an exogenous mechanical stimulus.

We also studied how local changes in the substrate stiffness affect the propagation of forces in the epithelium. Stiffness interfaces have been shown to affect the integrity of endothelial monolayers and impact their behavior over a distance of more than a hundred micrometers [57]. Similarly, we found that the cell displacement was affected near p_0_ even if the stiffness interface was at a distance of 80 *µ*m. Interestingly, when the forces propagate from soft to stiff substrate, the more distant interfaces led to a higher substrate displacement near p_0_. The origin of this effect is unclear. The cortical and junction forces on the soft region between the stiffness interfaces and p_0_ were increased, indicating that they experience more strain than on a substrate with a uniform stiffness. This is because the cells on the stiff region were more difficult to displace due to their stronger binding to the substrate and the stiffer substrate itself. A similar effect occurred for the force propagation from stiff to soft since the cells on the soft region were easier to move, meaning that more of the strain could be transmitted to them. Thus, this led to decreased cortical and junction forces in the stiff region.

The predicted peaks in the focal adhesion forces near the interface gradients with the increases in stiffnesses were a combination of two factors. First, the cells on the stiff substrate near the interface were subjected to a larger apical displacement from their close neighbors from the soft side. Second, the focal adhesion strength of these cells was higher due to the stiffer substrate under them. This combination thus led to the high relative increase in the focal adhesion forces. The situation was similar when the forces were transmitted from the stiff to the soft substrate. In this case, the cells on the soft side of the interface sensed smaller focal adhesion forces compared to the stiff uniform substrate due to the lower focal adhesion strength. However, since the cell displacement was already diminished at more distant interfaces, this effect was not as visible. Furthermore, we observed only minor differences in the cell displacement or forces in relation to the slope of the increase or decrease in stiffness. Therefore, whether the change in stiffness occurs within 2 or 50 *µ*m, the main factors that affected the cells were the change in stiffness and its distance.

We also studied how the substrate stiffness affects the small local changes in cell shape by implementing an optogenetic control of the myosin activation into our computational model. The results suggested that the substrate stiffness has only a minor effect on the small changes in the cell-cell junction elastic behavior. This can be explained by the cellular apico-basal connections, as small morphological changes in the apical side were not greatly restricted by the substrate-binding in the basal side of the cells. The observed difference in the relative substrate and cell displacement between the optogenetic and micromanipulation simulations show that the displacement of the apical side of the cells has to be extensive enough to visibly deform the substrate due to the compliance provided by the cells’ apico-basal axis.

The factors affecting the displacement and deformation of cells following some external force can be summarized as follows (Fig. 7). First, the stiffness of the substrate to which the cell is attached – together with the strength of this attachment – modulates the apical displacement. The second factor is the ability of the cells farther away from the external force to displace or deform. The cells move easily on a soft uniform substrate, as do the cells farther away. Therefore, the cell deformations are limited as it is easier to transmit the strain to the neighboring cells. On the other hand, the cell positions are more fixed on a stiffer substrate, making their movement more difficult, which leads to increased cell deformation. Furthermore, for example in the case with a stiffness increase opposite to the incoming force, the displacement of a cell in the soft region is limited by the less movable cells in the stiff region, leading to the more apical deformation. This further indicates that the information of the limited displacement farther away is transmitted as forces between the cells.

**Fig 7.**
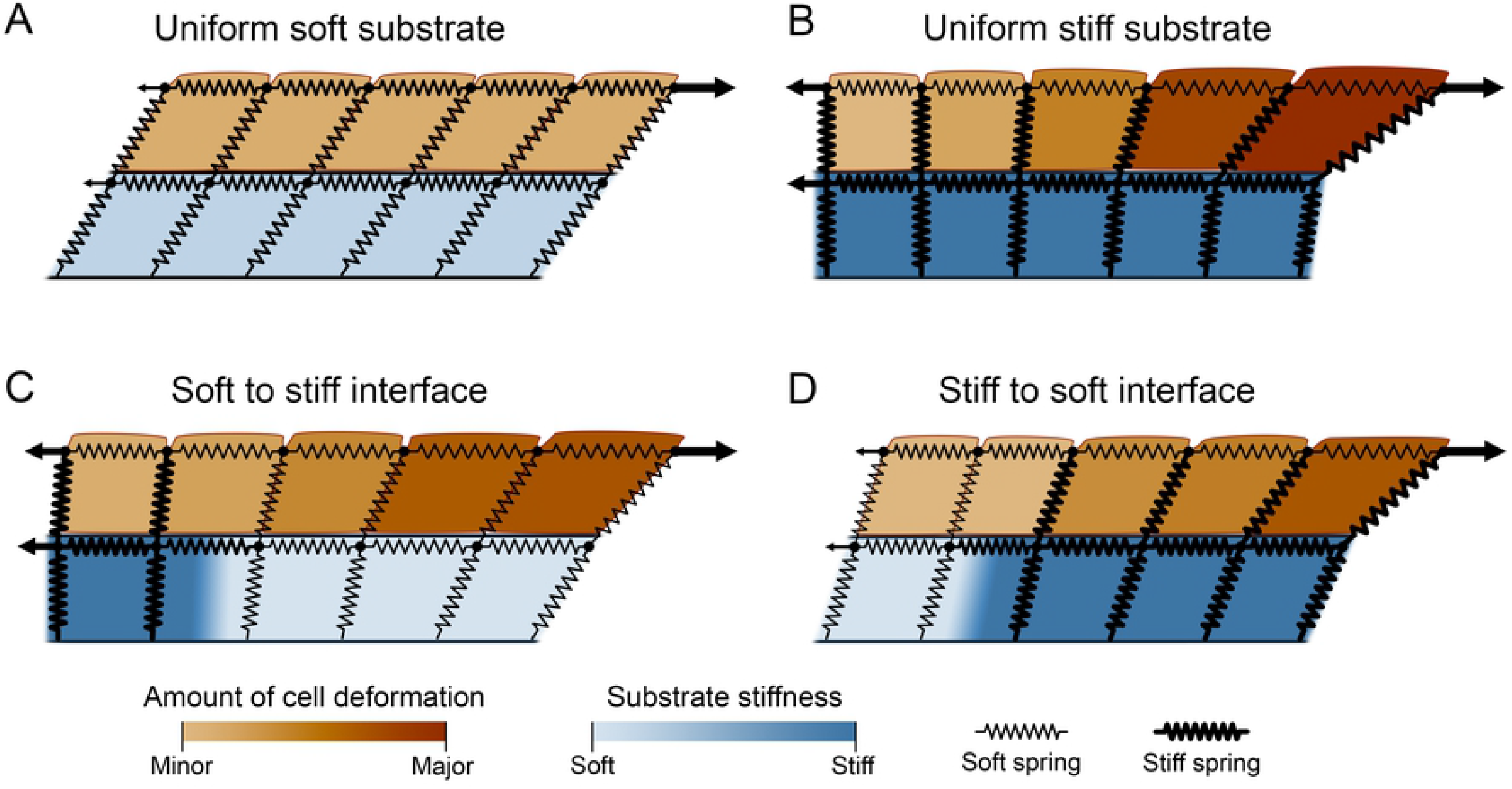
A 2D crosscut abstraction of the effect of substrate stiffness and stiffness interfaces on cell displacement and deformation upon mechanical stimulus. (A) On a uniform soft substrate, the cells are easily moved and do not substantially deform since the neighboring cells can also be readily moved. (B) A uniform stiff substrate is challenging to deform, leading to limited cell displacement, that is further limited by the stronger focal adhesion forces compared to the soft substrate. This leads to large deformation of the cells due to the limited ability of the neighboring cells to move. (C) When there is an increase in stiffness at a distance, the movement of cells on the stiff side of the stiffness interface is suppressed by the limited movement of the cells. This leads to more deformation of the cells on soft side of the interface compared to the uniform soft substrate. (D) With a decrease in stiffness at a distance, the cells on the soft substrate can readily move, enabling large cell movement also on the stiff substrate and thus less cell deformation compared to the uniform stiff substrate. The lengths in the figures are not to scale.

This communication of the forces and the information of the mechanical properties of the substrate is highly dependent on the cells’ resistance against deformation. If an external force can readily deform the cell’s apical surface, no force is left to transmit to the neighboring cells. Tension in the cytoskeleton has been shown to enable longer distance mechanical communication within cells compared to homogeneous solids [32, 58, 59], but it seems to be important also in the long-range mechanical signaling within an epithelial monolayer. This same effect is seen with fibrous substrates since separate cells have been shown to be able to communicate via the substrate over long distances [60–62].

The developed computational model forms a platform to complement the existing cell-based methods, by, to our knowledge, for the first time describing in detail the mechanics of the epithelial monolayer in combination with those of the underlying substrate. Compared to the common vertex model, our approach describes the cells and their interactions in more detail while being computationally considerably heavier. Furthermore, the model provides an additional level of complexity and dynamics compared to the previous closed-polygon-based models, e.g., Tamulonis et al. [48], since the cells are allowed to divide during growth and change their size and perimeter, and the junctions between the cells are allowed to remodel. A similar model to ours was recently published [63], building on the work by Tamulonis and coworkers with more dynamic cell functions and properties. However, like many other computational cell-based epithelial models, this model excluded the description of a deformable substrate under the epithelial monolayer. The few models that describe the substrate have not considered the effect of its mechanical properties in relation to the epithelial mechanics [42, 43]. In addition to the simulations presented here, the developed computational platform enables the description of further typical mechanical experiments conducted with epithelia, e.g., lateral substrate compression or stretching. Furthermore, we developed a graphical user interface for the simulations and data analysis to improve the platform’s usability.

While our modeling approach generally describes the system formed by the epithelial monolayer and the substrate well, there are limitations. First, the model cannot correctly describe the force propagation perpendicular to the micromanipulation on soft substrates, which can be explained by the rotation of the cell-cell junction interactions in relation to the cell membranes. Secondly, based on the slightly higher cell displacement at longer distances with the stiffer substrate predicted by the model compared to the experimental data, it seems that the elastic springs’ ability to describe the cell mechanics might be limited to cases with smaller strains. Furthermore, the description of the focal adhesion forces is challenging since they included both the focal adhesions themselves as well as the stiffness of the cell in the apico-basal axis. Separating these two components into their own forces could better describe the mechanics during the micromanipulation.

In summary, results from our *in vitro* cell model and computational simulations suggest that the mechanical properties of the substrate have a significant effect on the distance over which forces are transmitted in the epithelium. Furthermore, we found that the cells can communicate information of the substrate stiffness over long distances based on their ability to resist deformations. This indicates that, for example, the increased ECM stiffness in tumors can affect the mechanical signaling also outside the tumor itself. However, further studies are needed to better understand the role of each component in this phenomenon. The computational cell-based model presented here forms a valuable platform for futures studies on epithelial mechanics. In the future, the model would benefit from adding the tension remodeling described by Staddon et al. [24] and the inclusion of the cell nuclei. The latter would also allow the study of the forces felt by the nucleus and thus their possible role in regulating gene expression [29, 49, 64]. Furthermore, since the more fibrous nature of the natural ECM has been shown to transmit forces over longer distances [60–62], it would be interesting to study the ability of a fibrous substrate to propagate strain in the epithelial monolayer.

## Materials and methods

### Cell maintenance and establishment of MDCK mEmerald-Occludin-expressing cells

We used MDCK II (ATCC CCL-34) cells as an *in vitro* epithelial model tissue. The cells were cultivated in standard conditions in a humidified cell incubator (+37 °C, 5 % CO_2_) and maintained in Modified Eagle’s medium (#51200046, Thermo Fisher Scientific, Waltham, MA, USA) supplemented with 1 % (vol/vol) antibiotic (#15140122, Thermo Fisher Scientific) and 10 % fetal bovine serum (#10500064, Thermo Fisher Scientific). The MDCK cells used in the micromanipulation experiments stably expressed mEmerald-Occludin to highlight the cell-cell junctions with fluorescence. mEmerald-Occludin was a gift from Michael Davidson (Addgene plasmid #54212; http://n2t.net/addgene:54212; RRID: Addgene 54212). The MDCK mEmeral-Occludin cell line was established by first transfecting the MDCK cells with the mEmerald-Occludin plasmid using Neon Transfection system (Thermo Fisher, Waltham, USA). One day after the transfection, we started the positive cell selection with a medium where we replaced P/S with 0.75 mg/ml G418 antibiotic (#Gnl-41-01, Thermo Fisher Scientific). We picked positive colonies approximately two weeks later using a fluorescent microscope situated in the sterile cell culture hood. The MDCK mEmerald-Occludin cells were maintained in Modified Eagle’s medium (#51200046, Thermo Fisher Scientific, Waltham, MA, USA) supplemented with 0.25 mg/ml G418 antibiotic (#Gnl-41-01, Thermo Fisher Scientific) and 10 % fetal bovine serum (#10500064, Thermo Fisher Scientific).

### Polyacrylamide hydrogels and cell culturing

The polyacrylamide hydrogels were cast on 18 × 18 mm glass coverslips. First, coverslips were cleaned by immersing them in 2 % Helmanex for 1 h in +60 °C, followed by washes with excess water and ethanol. The coverslips were then let to dry in a fume hood or dried with a nitrogen stream. The cleaned coverslips were stored in a desiccator.

Before gel casting, the surfaces of the coverslips were amino-modified with 3-(Trimethoxysilyl)propyl methacrylate (#M6514, Sigma-Aldrich, Saint-Louis, USA) to allow firm gel attachment. The 3-(Trimethoxysilyl)propyl methacrylate and glacial acetic acid were mixed with 95 % ethanol yielding final concentrations of 0.3 % (vol/vol) and 5 % (vol/vol), respectively. The solution was let to react with a glass coverslip for 3 min at RT. Next, the coverslips were washed with excess ethanol and air-dried in a fume hood. The activated coverslips were stored in a desiccator.

The different gel rigidities were achieved by mixing different ratios of gel precursors acrylamide (AA) (stock 40 %, #1610140, Bio-Rad Laboratories, Hercules, USA) and bis-acrylamide (Bis) (stock 2 %, #1610142, Bio-Rad Laboratories, Hercules, USA) with PBS in 15 ml falcon tube [65]. The following mixing ratios were used: for 1.1 kPa gel final concentrations of AA and Bis were 3 % and 0.10 %, respectively; for 4.5 kPa 5 % and 0.15 %; for 11 kPa 10 % and 0.10 %; and for 35 kPa 10 % and 0.30 %. The gel precursor solution was then degassed with a vacuum. Next, 2 ml of this solution was pipetted into a new 15 ml falcon tube, and 2 % (vol/vol) fluorescent beads (0.2 *µ*m diameter, red fluorescent, #F8810, Thermo-Fisher, Waltham, USA) were added and mixed without bubble formation.

The gel polymerization was initiated by adding TEMED (#1610800, Bio-Rad Laboratories, Hercules, USA) and APS (10 % (weight/vol) stock solution in PBS, #A3678-100G, Merck, Kenilworth, USA) to a concentration of 0.2 % (vol/vol) and 1 % (vol/vol). The gel was mixed by tilting the tube 3–5 times, and immediately afterward, 13 *µ*l of gel solution was pipetted on an activated coverslip. Next, 13 mm cleaned but unactivated coverslip was carefully placed on top, sandwiching the polymerizing gel between the two coverslips. Gels were allowed to polymerize for 45 min in a moist chamber at RT. After polymerization, the gel-coverslip sandwiches were placed on 6-well plates, immersed in PBS, and kept o/n at +4 °C. On the following day, the 13 mm coverslips were carefully removed using a sharp scalpel, yielding approximately 100 *µ*m thick PAA gels on 18 *×* 18 mm coverslips.

Finally, the gels were coated with collagen-I to facilitate cell adhesion and growth. The coating was conducted by using 3,4-Dihydroxy-L-phenylalanine (L-DOPA) (#D9628, Sigma-Aldrich) according to Wouters et al. [66]. L-DOPA was dissolved in the dark to 10 mM Tris buffer, pH 10, with a final concentration of 2 mg/ml. The gel samples were incubated with L-DOPA solution for 30 min at RT in the dark. Next, the samples were washed twice with PBS and collagen-I in concentration of 50 *µ*g/ml in PBS was added on top of the gel and incubated for 1 h at RT. Finally, the cells were washed twice with PBS, and cell seeding was conducted immediately.

MDCK II cells stably expressing mEmerald-Occludin were maintained in 75 cm^2^ cell culture flasks. The protein-coated gels were placed on a sterile 6-well plates with PBS and sterilized in the laminar under UV light for 15 min. The cells were trypsinated and suspended into 10 ml of cell culture medium, and 100 *µ*l of the cell suspension was then pipetted on each gel, and 2 ml of medium was added to the well. Cells were cultured for 7 d prior to the micromanipulation experiments.

### Imaging and micromanipulation

We imaged the epithelial mechanics during micromanipulation using Nikon FN1 upright microscope (Nikon Europe BV, Amsterdam, Netherlands) with CFI Apo 40x/0.8 water-dipping objective. The mEmerald-Occludin was excited with 470 nm LED and beads with 580 nm LED from pE-4000 light source (CoolLED Ltd., Andover, UK). The system was equipped with W-VIEW GEMINI image splitting optics (Hamamatsu, Sunayama-cho, Japan), allowing simultaneous capturing of mEmerald-Occludin and fluorescent bead channels. The camera used in imaging was sCMOS ORCA-Flash 4.0 v2 (Hamamatsu, Sunayama-cho, Japan), which yielded an image pixel size of 330 nm. The used exposure time was 50 ms. During timelapse imaging, the frame rate was 13.4 frames per second.

Micromanipulation was conducted using uMp-3 triple-axis micromanipulator with uMp-TSC controller and uMp-RW3 rotary wheel interface (Sensapex, Oulu, Finland). The cells were manipulated by using a glass micropipette, similar to those used in patch-clamp recordings. The pipettes were constructed with P-1000 micropipette puller (Sutter Instruments, Novato, USA) and afterward closed with micro forge MF-830 (Narishige, Tokyo, Japan).

In the epithelium micromanipulation, the micropipette was brought in contact with a cell. The contact was visible in the microscopy images as a small indentation of the cell membrane. We started timelapse imaging and subsequently moved the micropipette 30 *µ*m with a speed of 30 *µ*m/s perpendicular to the pipette orientation. The movement was controlled via uMx Software (Sensapex, Oulu, Finland) using its macro commands. This yielded rapid mechanical manipulation of the cell and a movement of approximately 15 *µ*m of its center.

### Data analysis

The experimental imaging data before and after the micromanipulation pipette movement was initially segmented using the Trainable Weka Segmentation [67] plugin of ImageJ Fiji [68]. We randomly selected six images from the imaging data set to train the classifier to segment the cells based on the mEmerald-Occludin data to obtain the cell boundaries. Next, the probability maps were converted to binary masks using Find maxima and then skeletonized. We manually fixed any errors in the skeleton images based on comparison with the original images. Finally, BioVoxxel Toolbox’s Extended Particle Analyzer [69] was used to analyze the final segmented binary images. We tracked the movement of the cell centers between the segmented images before and after the pipette movements using a custom, semi-automated MATLAB script (R2020b, The MathWorks Inc., Natick, Massachusetts). The movement data was then used to interpolate the cell movement in relation to the original pipette position to obtain a cell movement map. Finally, we averaged the movement maps over the data from each gel stiffness.

We analyzed the gel deformation based on the fluorescent microbead data using Fiji’s particle image velocimetry (PIV) analysis plugin between the images before and after the micromanipulation. Similar to the cell data, the gel deformation maps were centered on the original pipette position and averaged over the same stiffnesses.

### Computational modeling

A detailed description of the model, the fitting, and the simulations are available in S2 Text. In our model, the epithelium was described as a two-dimensional monolayer, with each cell represented by a closed polygon (Fig. 8A). The model was based mainly on the boundary-based model by Tamulonis et al. [48] but borrowed features from the vertex models [40–42]. Cell structures and processes were incorporated into the model as forces affecting the polygon vertices. These include cortical actomyosin, cell-cell junction dynamics, intracellular pressure, cell division, focal adhesions, and membrane elasticity. Some of these forces are depicted in Fig. 8B. Furthermore, the cortical dynamics included the actomyosin prestress, described by a constant force component and a perimeter-dependent tension component. The number of the cell vertices was not static: new vertices were added to divide long membrane sections, and vertices were removed if a section between two vertices became too short. In addition, the cell-cell junctions were dynamic and constantly remodeled during the simulation.

**Fig 8.**
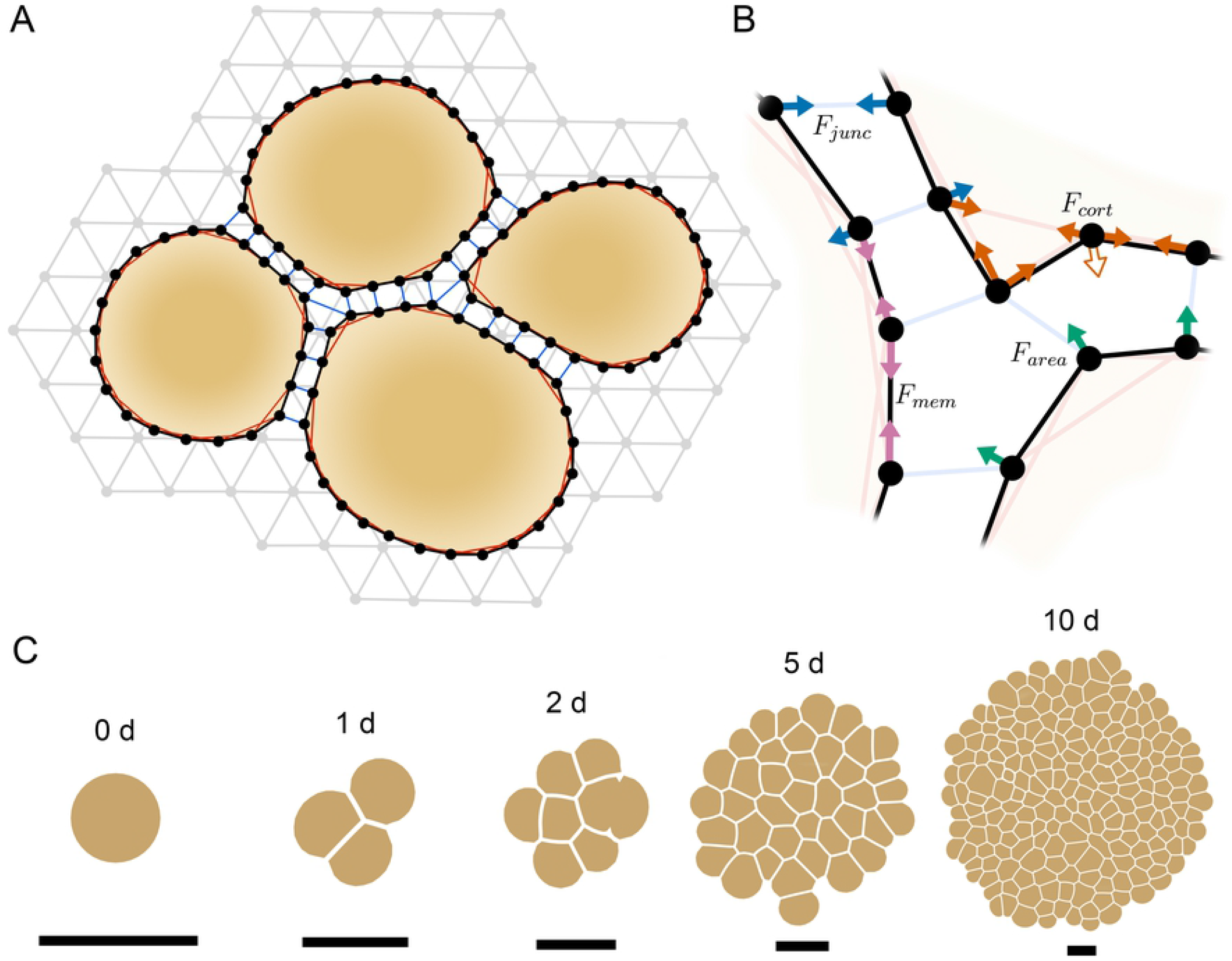
Depiction of the model. (A) Basic structure of the model. The cells were described by closed polygons and the cell structures and processes were included as forces affecting the polygon vertices. The cell-cell junctions and cortical actomyosin are depicted. The substrate was described by a triangular grid of points whose internal mechanics were defined by the forces between the grid points. The cell vertices were connected to the substrate via focal adhesion connections. (B) Example of forces that determine the cell vertex movements: cell-cell junction forces (F_*junc*_), cortical forces (F_*cort*_), membrane forces (F_*mem*_), and intracellular pressure or area force (F_*area*_). An additional cortical force component is added to the concave vertices, since the cortical link between its neighboring vertices runs behind it pushing it outwards (unfilled arrow). Note that the forces are calculated for every vertex but for simplicity, all forces are not shown for all vertices here. (C) Time series showing the growth of epithelial cell cluster from a single cell over a period of 10 days.

The top surface of the underlying substrate was represented by a two-dimensional triangular grid of points (Fig. **??**). As with the cells, the substrate mechanics were represented by forces acting on the grid points. The forces were related to the internal mechanics of the substrate as well as to the focal adhesions.

Equation of motion was used to evolve the model system during the simulation. The system was assumed to be overdamped, enabling the omission of inertial effects. This simplification is commonly done as the importance of inertia is small in biological systems [40, 41, 70]. The movement of cell vertex *i* and substrate point *m* were calculated as

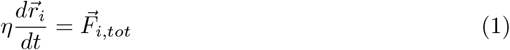

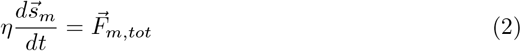

where η is the dampening coefficient (kg s^−1^), r_*i*_ is the position of the cell vertex *i* (m), s_*m*_ is the position of the substrate point *m* (m), t is time (s), and F_*i,tot*_ is the total force acting on cell vertex *i* (N) and F_*m,tot*_ that on the substrate point *m* (N). The total force for each cell vertex *i* was calculated as the sum of these component forces:

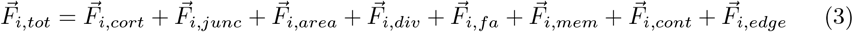

where 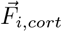 is the cortical actomyosin force, 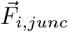 the cell-cell junction force, 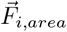 the area force that describes the internal pressure, 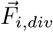 the division force, 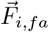 the focal adhesion force, 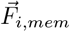 the membrane force, 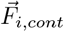 the contact force, and 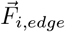 is the edge force. The last two forces had an auxiliary role: the contact force described contact between cells and prevented cell overlap, and the edge force described the continuity of the epithelium outside the simulated area.

The substrate mechanics were divided into three forces: a central force between neighboring points, a repulsive force between a point and the connection between two of its neighbors, and a restorative force that sought to move a point to its original location. The second force was included to prevent the collapse of the substrate during large deformations [71], and the third to describe the fact that the substrate was attached to rigid glass at its bottom surface in our experiments. Furthermore, a fourth force component was included to depict the cell-substrate connection via the focal adhesions. Now, the total force affecting each substrate point was calculated as

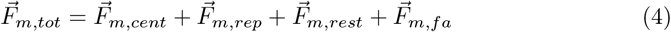

where 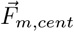 is the central force between closest neighboring points, 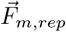 is the repulsive force to prevent material collapse, 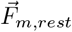 is a restorative force, and 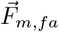 is the force from the focal adhesions.

The model was used to simulate epithelial growth and the tissue response to two different mechanical stimuli: 1) pointlike micromanipulation in a short time scale and 2) a local increase in actomyosin tension by optogenetics over a longer time scale.

We used the model to grow epithelia from a single cell (Fig. 8C) to produce epithelium of sufficient size without the substrate. The randomness in the tissue was produced by normally distributed times between divisions and cell area distribution based on our *in vitro* MDCK cell data. The size of the grown epithelium was chosen based on the assumed effect of each mechanical stimulus to minimize the impact of the tissue edges. Following the growth, the epithelia were given time to relax without division to remove any stresses. Next, the grown epithelia were placed on the substrate, and the focal adhesions were defined between the two.

Corresponding to our micromanipulation experiments, we moved a single cell by an external force with a known speed over a distance. Since we wanted to describe the elastic behavior, we prohibited any changes in the number of cell vertices and cell-cell junctions in these simulations, justified by the short time scale of these measurements. The values of the model parameters governing the cell mechanics were fitted using our *in vitro* micromanipulation data with the uniform 1.1, 4.5, and 11-kPa substrates by iteratively changing the parameter values and comparing the cell center and substrate displacements between the experimental data and simulations results. The fitted model was then used to study the force propagation on the uniform substrates and those with stiffness interfaces and gradients. The interfaces and the gradients were defined along the direction of the virtual pipette movement and characterized by the gradient slope and distance from the initial pipette position.

In the optogenetic activation simulations, the contractility of the cortex in a section between two randomly chosen cells was increased to describe the experimental myosin activation. This was done by increasing the value of cortical tension constants for the cortical forces within the activation region. The parameters for these simulations were obtained from Staddon et al. [24] and by fitting our model to their data.

The model was solved using either 2nd or 4th order Runge-Kutta methods with variable time steps. During the growth simulations when the substrate was excluded, 2nd order Runge-Kutta was used since it was sufficiently accurate. These simulations also omitted the focal adhesion and the cell edge forces. During the simulations that included the substrate, 4th order Runge-Kutta was used to evolve the system.

The model is implemented in MATLAB, where we also created a graphical user interface for the model platform. The model code is available in GitHub (https://github.com/atervn/epimech).

## Supporting information

### S1 Appendix. Supporting figures

Figures showing the cell and substrate displacement data and the force data from which the stiffness interface gradient results were calculated (Figs. A and C). Figures show the cell and substrate displacement data and the force data, and the difference results for the sharp and shallow stiffness gradients (Figs. B and D). An example of the forces affecting the cell vertices and substrate points following 30-*µ*m micromanipulation of a single cell (Fig. E).

### S2 Text. Description of the EpiMech model

A detailed description of the model, the included cell components and processes, the forces, the simulations, and the parameter values.

### S3 Video. Experimental micromanipulation

Representative examples of the micromanipulation experiments showing both the cell boundaries and the fluorescent microbeads of the hydrogel substrates for the stiffnesses 1.1, 4.5, 11, and 35 kPa.

### S4 Video. Simulated micromanipulation

Representative examples of the micromanipulation simulations showing both the cell boundaries and the substrate points for the stiffnesses 1.1, 4.5, and 11 kPa. The purple cross represents the micromanipulator position.

### S5 Video. Simulated growth

Representative example of an epithelium growth simulation from a single cell over 14 hours.

### S6 Video. Optogenetics simulation

Representative example of an optogenetic activation simulation showing both the cell boundaries and the substrate points. The simulation was run with a substrate with a stiffness of 1.1 kPa. The purple region shows the activated area during the light activation from 2 to 22 minutes.

## Acknowledgments

The authors acknowledge the Biocenter Finland (BF), Tampere Imaging Facility (TIF), and Tampere facility of Electrophysiological Measurements for the service. Michael Davidson is kindly acknowledged for the mEmerald-Occludin plasmid (Addgene plasmid #54212).

